# Mice lacking NF-κB1 exhibit marked DNA damage responses and more severe gastric pathology in response to intraperitoneal tamoxifen administration

**DOI:** 10.1101/112656

**Authors:** Michael D Burkitt, Jonathan M Williams, Tristan Townsend, Rachael Hough, Carrie A Duckworth, D Mark Pritchard

## Abstract

**Background:** Tamoxifen (TAM) has recently been shown to cause acute gastric atrophy and metaplasia in mice. We have previously demonstrated that the outcome of *Helicobacter felis* infection, which induces similar gastric lesions in mice, is altered by deletion of specific NF-κB subunits. *Nfkb1*^-/-^ mice developed more severe gastric atrophy than wild-type (WT) mice 6 weeks after *H. felis* infection. In contrast, *Nfkb2*^-/-^ mice were protected from this pathology. We therefore hypothesized that gastric lesions induced by TAM may be similarly regulated by signaling via NF-κB subunits.

**Methods:** Groups of 5 female C57BL/6 (WT), *Nfkb1*^-/-^, *Nfkb2*^-/-^ and *c-Rel*^-/-^ mice were administered 150mg/kg TAM by IP injection. 72 hours later, gastric corpus tissues were taken for quantitative histological assessment. In addition, groups of 6 female WT and *Nfkb1*^-/-^ mice were exposed to 12Gy γ-irradiation. Gastric epithelial apoptosis was quantified 6 and 48 hours after irradiation.

**Results:** TAM induced gastric epithelial lesions in all strains of mice, but this was more severe in *Nfkb1*^-/-^ mice than WT mice. *Nfkb1*^-/-^ mice exhibited more severe parietal cell loss than WT mice, had increased gastric epithelial expression of Ki67 and had an exaggerated gastric epithelial DNA damage response as quantified by γH2AX. To determine investigate whether the difference in gastric epithelial DNA damage response of *Nfkb1*^-/-^ mice was unique to TAM induced DNA damage, or a generic consequence of DNA damage, we also assessed gastric epithelial apoptosis following γ-irradiation. 6 hours after γ-irradiation, gastric epithelial apoptosis was increased in the gastric corpus and antrum of *Nfkb1*^-/-^ mice.

**Conclusions:** NF-κB1 mediated signaling regulates the development of gastric mucosal pathology following TAM administration. This is associated with an exaggerated gastric epithelial DNA damage response. This aberrant response appears to reflect a more generic sensitization of the gastric mucosa of *Nfkb1*^-/-^ mice to DNA damage.

## Introduction

Gastric cancer in humans is strongly associated with gastric colonization with *Helicobacter pylori*^1^. This leads, in a minority of people, to a pre-neoplastic cascade of pathology which develops over several decades. This cascade is typified by gastric oxyntic gland atrophy, epithelial metaplasia, dysplasia and cancer^2^. These events can be modelled by infecting C57BL/6 mice with the related bacterium *H. felis.* In this model, advanced gastric pre-neoplasia develops over 12 months and, in some hands, malignancies have been reported in animals aged 13-15 months^3,4^.

The development of gastric pre-neoplasia in response to gastric *Helicobacter* infection occurs on the background of chronic inflammation of the gastric mucosa. The signaling pathways that regulate gastric epithelial remodeling in response to chronic inflammation are complex and have not been fully elucidated. However there is a wealth of evidence that signaling through NF-κB pathways is involved in the development of inflammation associated malignancies both in the stomach, and in other parts of the gastrointestinal tract^5^. The NF-κB proteins are a group five of structurally related transcription factors (RelA (p65), RelB, c-Rel, NF-κB1 (p50) and NF-κB2 (p52)) which influence transcription through the binding of homo- or hetero-dimers of themselves to chromatin. Activation of transcription through these proteins is tightly regulated, both through a network of upstream signaling kinases, and by complex post transcriptional modification of the NF-κB subunits. Two distinct NF-κB pathways are conventionally described: the classical (canonical) pathway is regulated by the activation of IKKβ, following stimuli including TNF binding to its receptor, and activation of toll-like receptors. The principal NF-κB dimers activated by this pathway are RelA/NF-κB1 dimers, and RelA/c-Rel dimers. In contrast, the alternative (non-canonical) NF-κB signaling pathway is regulated by the kinase NIK, and is characterized by transcriptional regulation by dimers of RelB/NF-κB2 (see Merga *et al* for a recent review of these pathways^5^).

We have previously reported that *H. felis* induced gastric pre-neoplasia is differentially regulated by signaling involving specific NF-κB sub-units. Mice lacking the NF-κB1 subunit developed more severe gastric atrophy after 6 weeks of infection, and more severe pre-neoplastic pathology when infected for 12 months. In contrast, *Nfkb2*^-/-^ mice were entirely protected from *H. felis* induced pathology, despite heavy colonization by these bacteria^6^. This has also been demonstrated in the context of acute inflammatory responses in the intestinal tract, where *Nfkb2*^-/-^ mice exposed to low dose lipopolysaccharide (LPS) systemically, were protected from pathological small intestinal villus tip epithelial cell shedding and apoptosis whereas *Nfkb1*^-/-^ mice demonstrated more severe lesions than wild-type mice^7^.

It has recently been reported that intraperitoneal administration of tamoxifen to mice induces an acute gastric corpus metaplasia, characterized by the loss of parietal cells, metaplastic changes originating in the chief cells at the base of the gastric glands, and increased epithelial cell proliferation^8^. These findings have been shown to be reproducible in other laboratories^9^, to be estrogen independent and reversible on discontinuation of tamoxifen. Unlike *H. felis* induced gastric pre-neoplasia, this pathology occurs with relatively little associated inflammation in the gastric mucosa^10^; this model therefore offers an opportunity to investigate gastric epithelial remodeling in the absence of a chronic inflammatory stimulus.

We have utilized this model to characterize whether the regulation of gastric pre-neoplasia by mice lacking specific NF-κB sub-units is a generic response to gastric epithelial remodeling, or specific to the events induced by *H. felis* infection.

## Results

### Tamoxifen induced gastric corpus pathology is regulated by Nfkb1

To characterize whether deletion of specific NF-κB sub-units altered the severity of pathology induced by administration of tamoxifen, groups of at least 5, 12-week old, WT, *Nfkb1*^-/-^, *Nfkb2*^-/-^ and *c-Rel*^-/-^ female mice were either treated with tamoxifen, or vehicle and culled 72 hours later. Gastric pathology was scored using an established scoring system^11^. Vehicle treated mice of all genotypes exhibited minimal gastric lesions (Figures 1 and 2A). Following administration of tamoxifen morphological changes were seen in the corpus mucosa of all groups of mice. Tamoxifen treated WT mice had mean pathology scores of 2.4(+/- 0.51 SEM). *Nfkb2*^-/-^ and *c-Rel*^-/-^ mice exhibited similar lesion scores, whilst *Nfkb1*^-/-^ mice exhibited more severe gastric lesions with a mean score of 4.8(+/- 0.74, *p*<0.01 by 2-way ANOVA and Dunnett’s *post-hoc* test).

To quantify the degree of gastric atrophy induced by tamoxifen, we assessed the number of H^+^/K^+^ATPase expressing cells. Amongst untreated mice, no significant differences in the number or distribution of parietal cells were identified between the mice of different genotypes. 31.2% of cells in the gastric corpus of untreated WT mice expressed H^+^/K^+^ATPase (Figures 1 and 2B), and cells were distributed between cell positions 2 and 36 of the gastric gland, with peak prevalence at cell position 15 (Figure 2C).

**Figure 1:**
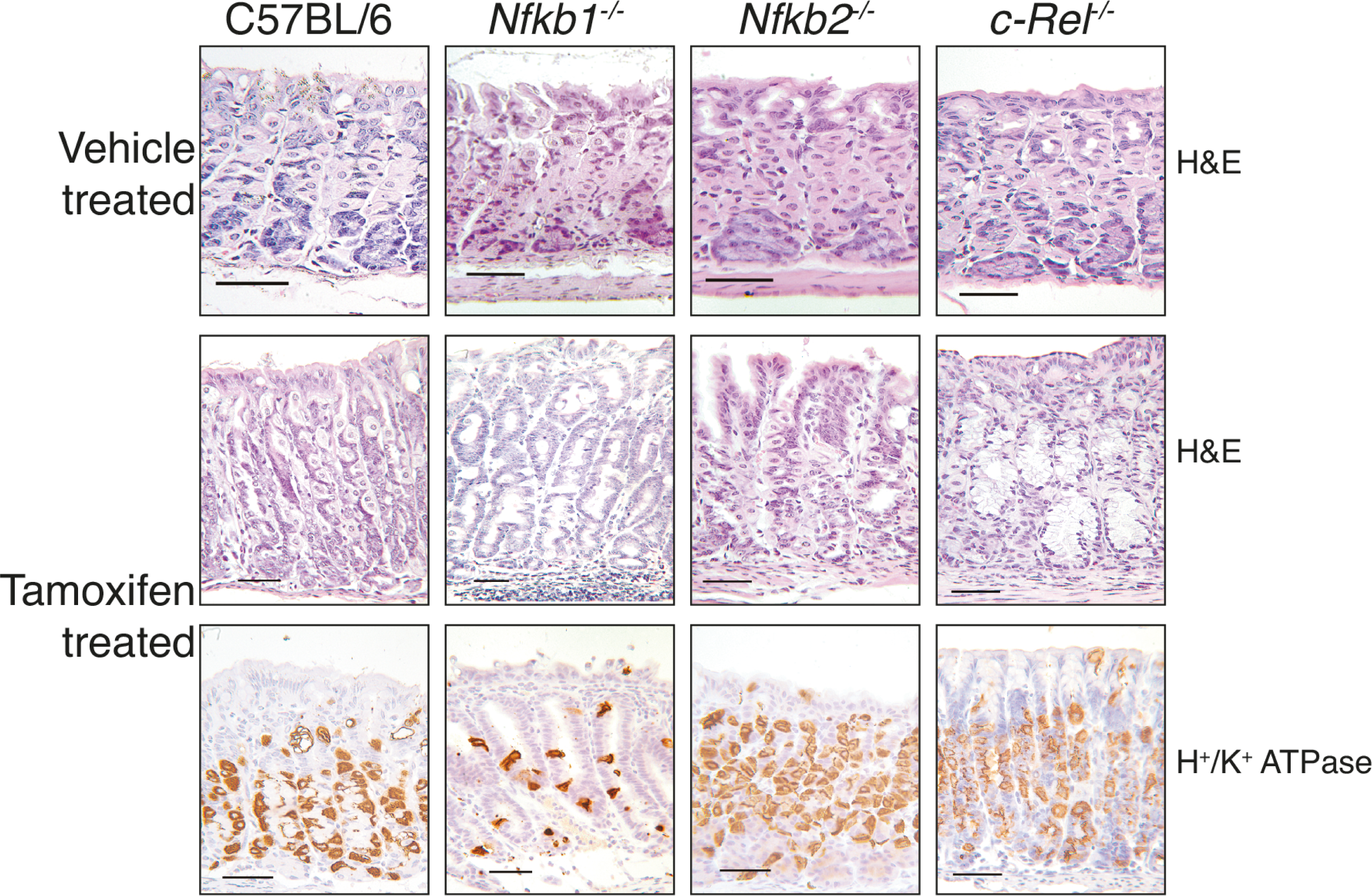
Representative photomicrographs of gastric corpus of WT, *Nfkb1*^-/-^, *Nfkb2*^-/-^ and *c-Rel*^-/-^ mice treated with vehicle or 150mg/kg tamoxifen. Sections stained with hematoxylin and eosin, or immunostained for expression of H^+^/K^+^ATPase. Scale bars 100μm.

**Figure 2:**
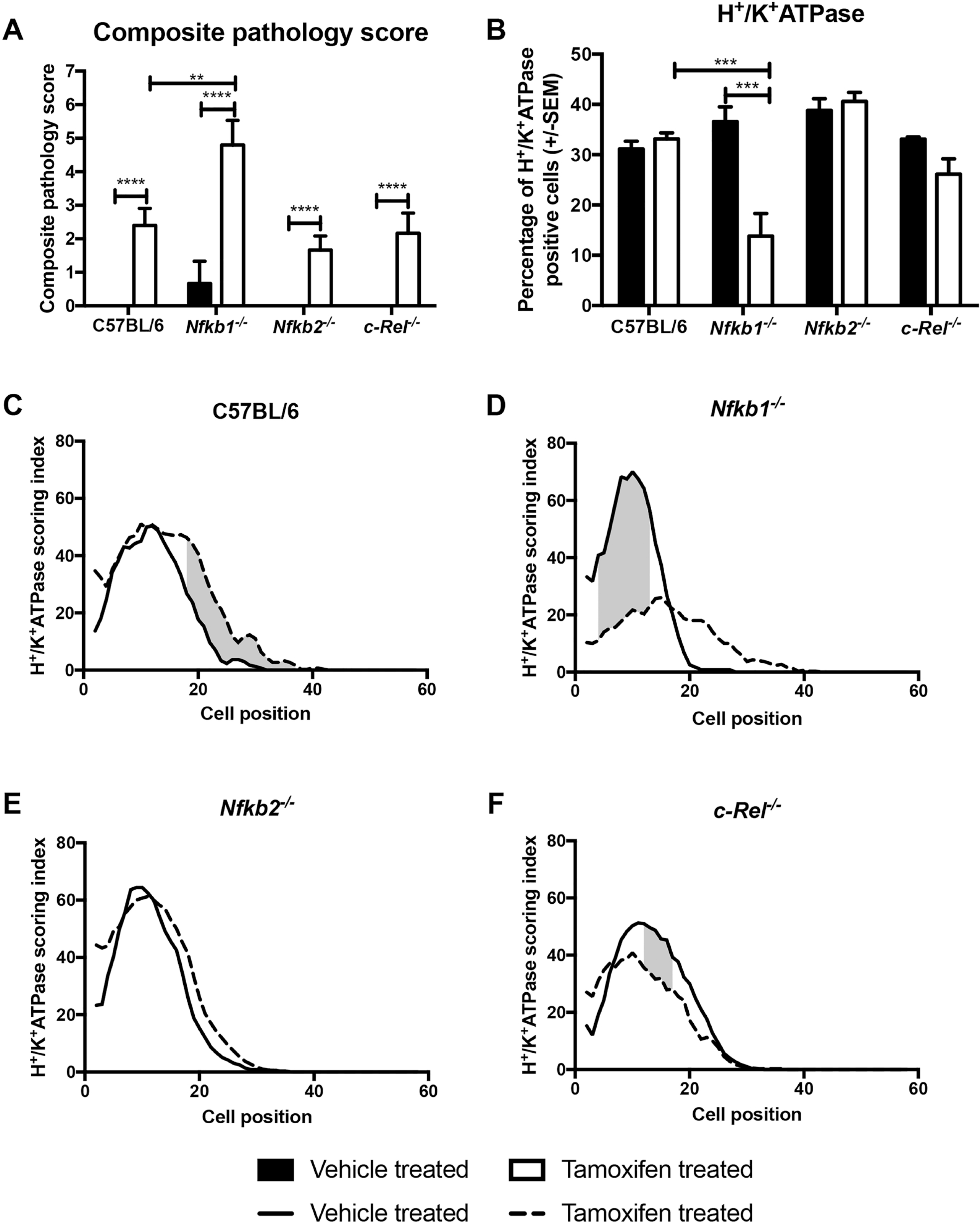
Tamoxifen induced epithelial lesions in WT, *Nfkb1*^-/-^, *Nfkb2*^-/-^ and *c-Rel*^-/-^ mice. A: visual analogue scoring of pathology identified in H+E stained sections from mice treated with vehicle or 150mg/kg tamoxifen. B: percentage of gastric corpus gland cells expressing H^+^/K^+^ATPase in mice treated with vehicle or 150mg/kg tamoxifen. For A and B statistically significant differences tested by 2-way ANOVA and Dunnett’s test. ** *p*<0.01, *** *p*<0.001, **** *p*<0.0001. C-F: H^+^/K^+^ATPase positive cells plotted by cell position in the gastric corpus hemiglands of WT (C), *Nfkb1*^-/-^ (D), *Nfkb2*^-/-^(E) and *c-Rel*^-/-^(F). Solid line represents distribution of cells in vehicle treated mice, broken line tamoxifen treated mice. Shaded area marks region of gland where distribution of H^+^/K^+^ATPase positive cells differs significantly in tamoxifen treated vs vehicle treated mice, *p*<0.05 by modified median test. N=5 for all experimental groups.

**Table 1:**
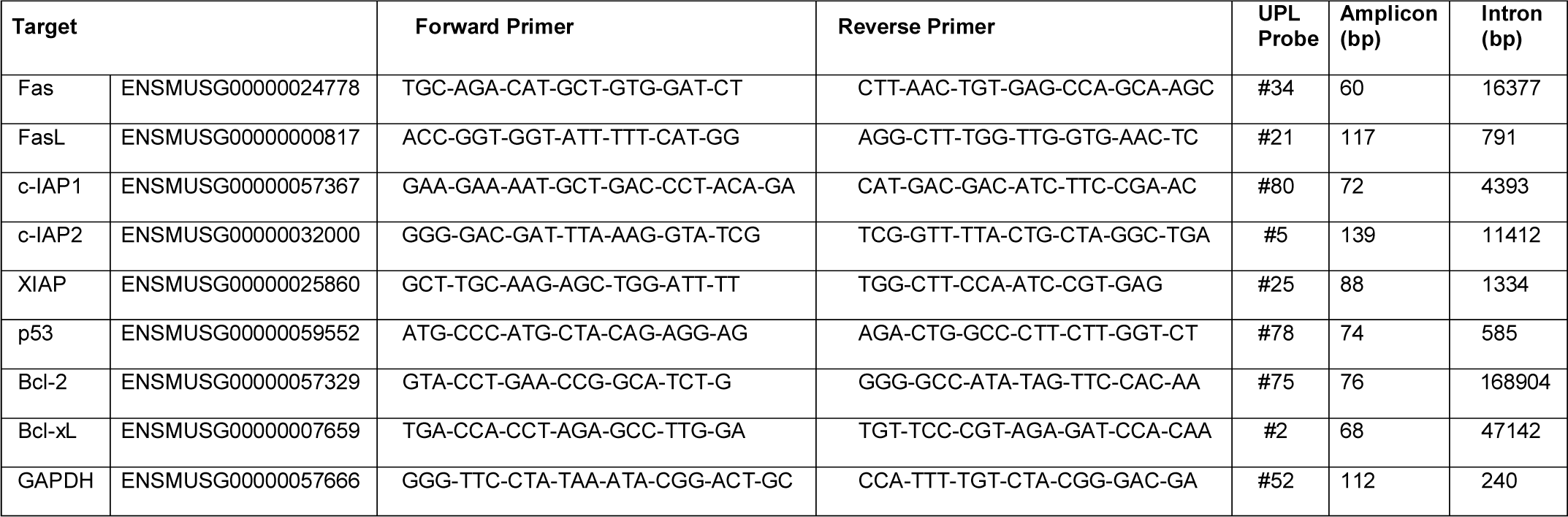
Primers, probes and amplicons for quantitative PCR assays.

Following administration of tamoxifen, the number of H^+^/K^+^ATPase expressing cells did not differ substantially in WT mice, but a shift in parietal cell distribution up the gland was noted (significantly higher numbers of parietal cells observed between cell positions 18 and 36 in tamoxifen treated mice, p<.05 by modified median testing, Figure 2C). In keeping with the visual analogue scoring of gastric lesions, similar numbers and distributions of H^+^/K^+^ATPase expressing cells were identified in the corpus of *Nfkb2*^-/-^ and *c-Rel*^-/-^ mice that had been exposed to tamoxifen (Figure 2B, E and F). In contrast, *Nfkb1*^-/-^ mice showed more marked changes in parietal cell distribution than their WT counterparts, with a 2.4-fold reduction in parietal cell number (*p*<.001 by 2-way ANOVA and Dunnett’s *post-hoc* analysis, Figure 2B), and a marked reduction in parietal cell abundance between cell positions 4 and 13 of gastric corpus glands (*p*<0.05 by modified median testing, Figure 2D). To validate this observation, and to determine whether the observed effects of tamoxifen were likely to be due to an estrogen receptor mediated effect, we performed quantitative real-time PCR on gastric mucosal samples from WT and *Nfkb1*^-/-^_mice that had, or had not been treated with tamoxifen. These assays demonstrated reduced transcription of ATP4A (encoding H^+^/K^+^ATPase)in tamoxifen treated WT and *Nfkb1*^-/-^ mice, but no differences in expression of the estrogen receptor regulated genes Erbb2 or Wnt5A (supplementary figure 1).

### Nfkbl mediated signaling regulates tamoxifen induced cell proliferation

To determine whether NF-κB signaling influenced gastric epithelial cell turnover following the administration of tamoxifen, we quantified cells expressing the S-phase marker Ki67, and those undergoing apoptosis by expression of cleaved-caspase 3.

In untreated mice, genotype did not significantly influence the number of cleaved-caspase 3 positive cells, which were relatively rare occurrences (1.4+/-0.68 SEM cells per high power (x40 objective) field; HPF). However there was a trend towards more apoptosis in both *Nfkb1*^-/-^ and *Nfkb2*^-/-^ mice. Administration of tamoxifen to WT mice increased the number of cleaved-caspase 3 positive cells 3.4-fold, but this did not reach statistical significance (Figure 3A and B). In both *Nfkb1*^-/-^ and *Nfkb2*^-/-^ mice tamoxifen induced apoptosis in a larger number of apoptotic cells than in WT mice (3.9- and 4.8-fold, *p*<0.05 and *p*<0.01 respectively).

**Figure 3:**
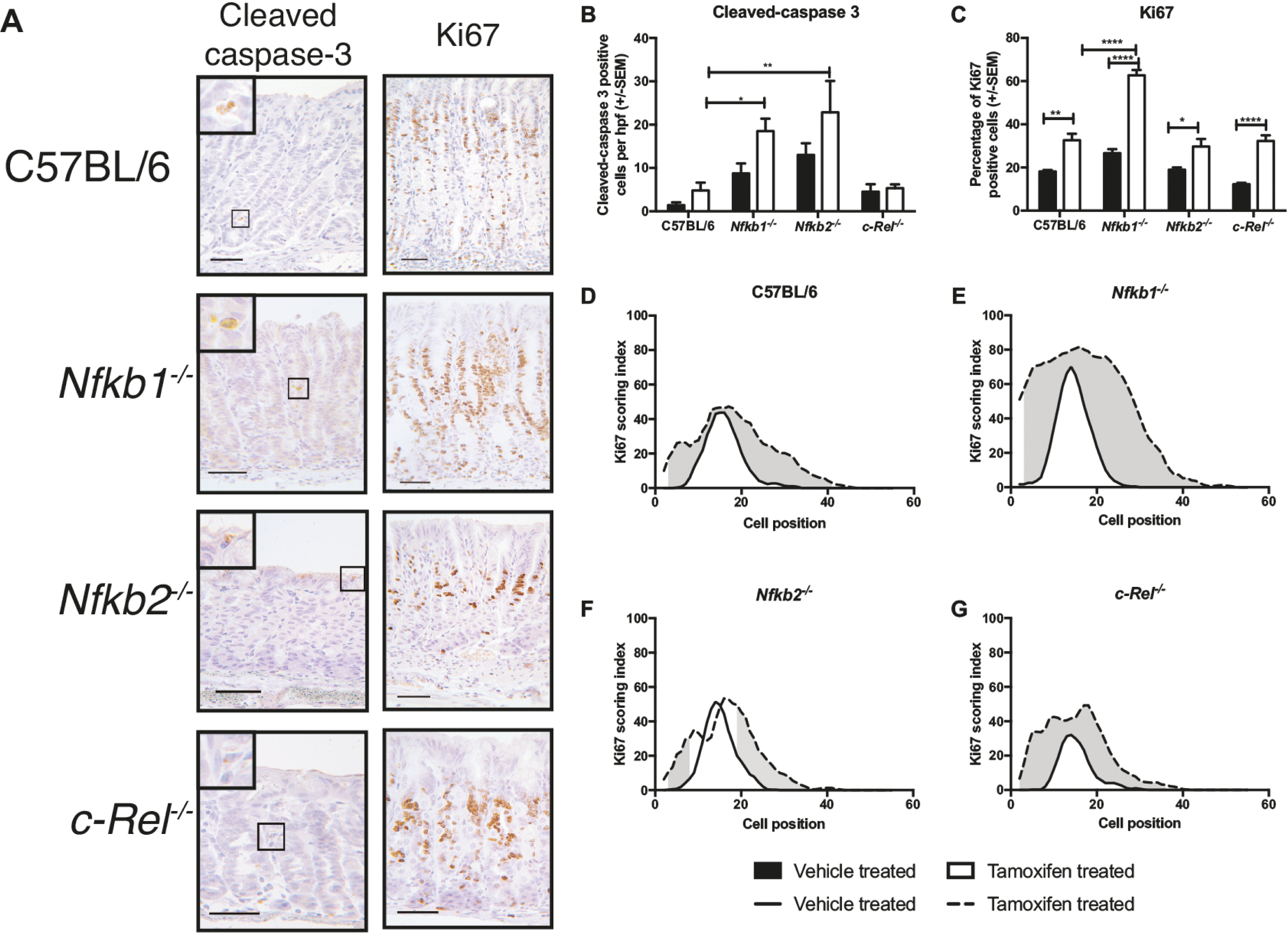
Gastric corpus epithelial cell turnover following tamoxifen administration in WT, *Nfkb1*^-/-^, *Nfkb2*^-/-^ and *c-Rel*^-/-^ mice. A: Representative photomicrographs of gastric corpus of WT, *Nfkb1*^-/-^, *Nfkb2*^-/-^ and *c-Rel*^-/-^ mice treated with 150mg/kg tamoxifen. Sections immunostained for expression of cleaved caspase-3 or Ki67. Scale bars 100μm. B: Number of cleaved-caspase 3 positive cells identified per high power field in mice treated with vehicle or 150mg/kg tamoxifen. C: percentage of gastric corpus gland cells expressing Ki67 in mice treated with vehicle or 150mg/kg tamoxifen. For B and C statistically significant differences tested by 2-way ANOVA and Dunnett’s test. * *p*<0.05, ** *p*<0.01, **** *p*<0.00001. C-F: Ki67 positive cells plotted by cell position in the gastric corpus hemiglands of WT (D), *Nfkb1*^-/-^ (E), *Nfkb2*^-/-^(F) and *c-Rel*^-/-^(G) mice. Solid line represents distribution of cells in vehicle treated mice, broken line tamoxifen treated mice. Shaded area marks region of gland where distribution of Ki 67 positive cells differs significantly in tamoxifen treated vs vehicle treated mice, *p*<0.05 by modified median test. N=5 for all experimental groups.

In untreated WT mice 18.1%(+/-0.7) of gastric corpus epithelial cells were in S-phase. They were distributed between cell positions 5 and 32, peak proliferative index was observed at cell positions 15 and 16. Proliferating cells were more abundant in untreated *Nfkb1*^-/-^ mice, compared to WT (26.6%+/-1.9, *p*<.05 by 2-way ANOVA and Dunnett’s post-hoc test); in this genotype, peak proliferative index was observed at cell position 14, and proliferating cells were observed between cell positions 2 and 29. Other untreated transgenic mice exhibited similar proliferation indices to WT mice.

Following tamoxifen treatment, the number of proliferating cells observed in WT mice increased 1.8-fold (*p*<0.01, Figures 3A, C and D). This increase was recapitulated in other genotypes of mice (Figures 3A, C, E, F and G), but was more pronounced in *Nfkb1*^-/-^ mice than other groups, where a 2.4-fold increase in proliferating cells was observed (*p*<0.0001 compared with tamoxifen treated WT mice, Figure 3C). In these animals proliferating cells were observed between cell positions 2 and 52, with peak proliferative index at cell position 16.

### Nfkb1 regulates tamoxifen induced DNA damage in the gastric epithelium

To investigate the mechanism underlying differences in sensitivity to tamoxifen induced gastric pathology we assessed whether there were any aberrant DNA damage responses. Whilst tamoxifen is best characterized as an anti-estrogen receptor drug, it is also a genotoxic agent. DNA damage induced by tamoxifen is associated with the formation of tamoxifen/DNA adducts, and with the induction of oxidative stress by tamoxifen metabolites^12^. This mechanism has been hypothesized to be of clinical relevance in patients treated with tamoxifen, who have an increased risk of some malignancies, including endometrial cancer^13^. We hypothesized therefore, that this effect may influence the degree of gastric pathology induced by tamoxifen.

To quantify this, we immunostained tissue sections using a phospho-specific antibody targeting γ-H2AX, which reflects a DNA damage response in the cell. In untreated gastric corpus mucosa, γ-H2AX was identified in 2.8% of WT epithelial cells. All untreated transgenic groups had similar percentages of γ-H2AX labelled cells to wild-type mice (Figure 4A-F). There was a small increase in abundance of γ-H2AX following tamoxifen treatment of WT mice (4.6%), but this did not reach statistical significance. A similar trend towards increased abundance of γ-H2AX was observed in *NfKb2*^-/-^ and c-*Rel*^-/-^ mice administered tamoxifen. In contrast, the percentage of γ-H2AX labelled cells increased 4.0-fold in *Nfkb1*^-/-^ mice administered tamoxifen (*p*<0.001, Figure 4B). Excess γ-H2AX expression occurred over a wide gland area in this strain (between cell positions 5 and 32 of the gastric corpus gland, Figure 4D). In tamoxifen treated *Nfkb1*^-/-^ mice, γ-H2AX expression was observed between cell positions 2 and 39, with peak γ-H2AX expression observed at cell position 20 (Figure 4D).

**Figure 4:**
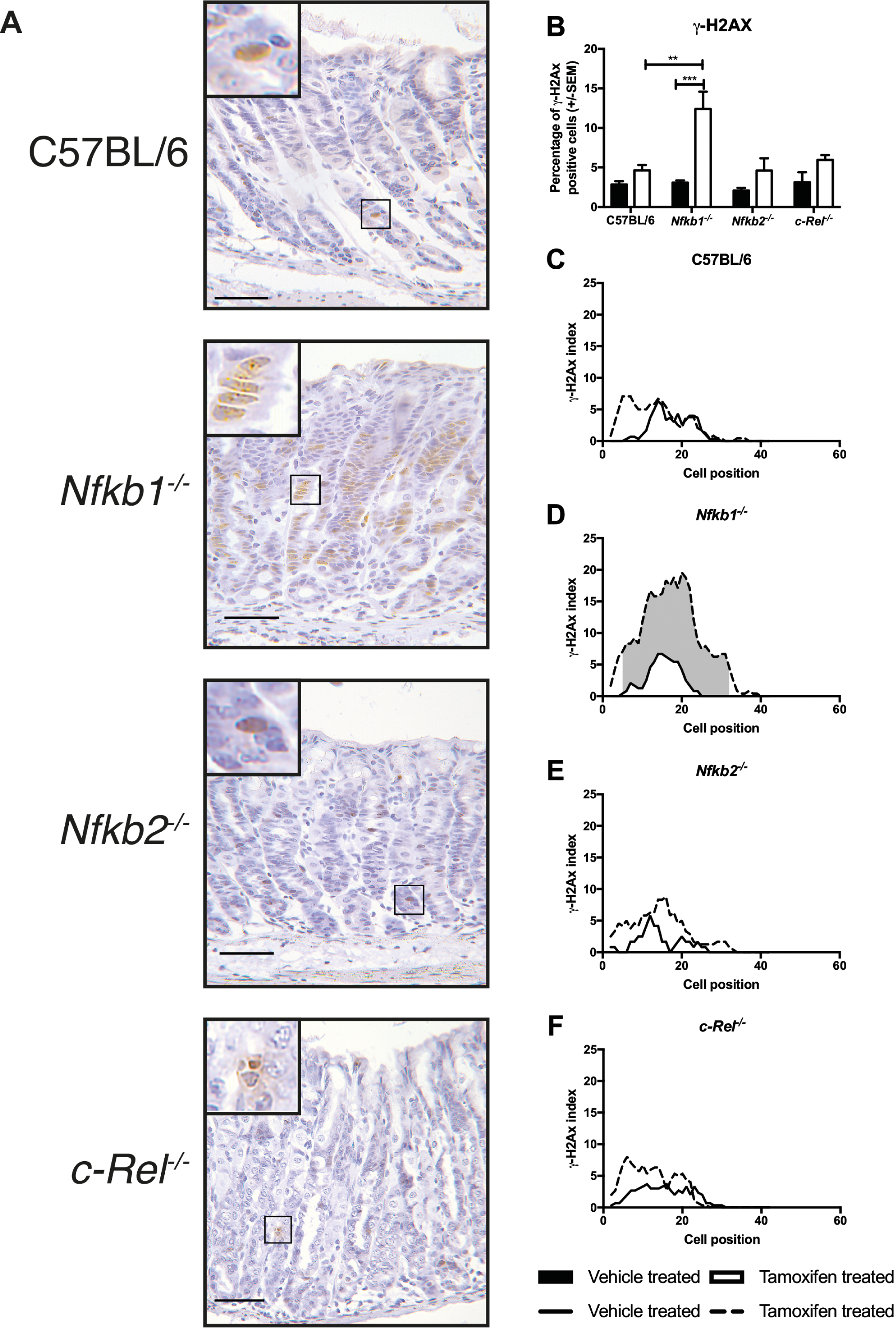
Gastric corpus expression of γ-H2AX following tamoxifen administration in WT, *Nfkb1*^-/-^, *Nfkb2*^-/-^ and *c-Rel*^-/-^ mice. A: Representative photomicrographs of gastric corpus of WT, *Nfkb1*^-/-^, *Nfkb2*^-/-^ and *c-Rel*^-/-^ mice treated with 150mg/kg tamoxifen. Sections immunostained for expression of γ-H2AX. Scale bars 100μm. B: percentage of gastric corpus gland cells expressing γ-H2AX in mice treated with vehicle or 150mg/kg tamoxifen. Statistically significant differences tested by 2-way ANOVA and Dunnett’s test. ** *p*<0.01, *** *p*<0.001. B-E: γ-H2AX positive cells plotted by cell position in the gastric corpus hemiglands of WT (C), *Nfkb1*^-/-^ (D), *Nfkb2*^-/-^ (E) and *c-Rel*^-/-^ (F) mice. Solid line represents distribution of cells in vehicle treated mice, broken line tamoxifen treated mice. Shaded area marks region of gland where distribution of γ-H2AX cells differs significantly in tamoxifen treated vs vehicle treated mice, *p*<0.05 by modified median test. N=5 for all groups.

Previous studies have suggested that the administration of tamoxifen specifically induced gastric parietal cell death. To characterize which cell types experienced DNA damage in our studies we performed co-immunofluorescence studies with immunostaining for both H^+^/K^+^ATPase and γ-H2AX. These assays demonstrated that whilst some γ-H2AX positive cells were parietal cells, other γ-H2AX positive cells did not express H^+^/K^+^ATPase (supplementary figure 2). Because of the differences in proliferation indices observed in *Nfkb1*^-/-^ mice following TAM administration, we also wanted to determine whether DNA damage, identified by γ-H2AX immunostaining, co-localised predominantly with proliferating cells. To do this we performed γ-H2AX/Ki67 co-immunostaining. This assay demonstrated that whilst a proportion of γ-H2AX stained cells were in S-phase following tamoxifen administration, the others did not appear to be proliferating (supplementary figure 3).

### Nfkb1^-/-^ mice also have aberrant responses to γ-irradiation induced DNA damage

We next wanted to characterize whether the differences identified in γ-H2AX expression in *Nfkb1*^-/-^ mice were due specifically to altered responses to the genotoxic stress induced by tamoxifen, or represented a more generic aberrant response to DNA damage. To address this, we quantified gastric epithelial cell turnover in both the gastric corpus and antrum at baseline, and 6 and 48 hours following 12Gy γ-irradiation, by cell positional scoring of morphologically apoptotic and mitotic cells. This dose and time-points of γ-irradiation have previously been demonstrated to be optimal for the assessment of DNA damage induced apoptosis in the stomach, and morphological assessment of H+E stained sections correlates well with other markers of apoptosis including cleaved-caspase 3 immunostaining and TUNEL staining^14^.

In untreated age matched, young adult WT mice, gastric antral glands were 19.1(+/-1.9) cells in length on average, whilst corpus glands were a mean 31.1(+/-1.7) cells long. Deletion of NF-κB subunits did not significantly alter the length of gastric corpus glands, whilst in the antrum, gastric gland hyperplasia to a mean gland length of 24.2 cells (+/- 2.6, *p*<.01) was observed in *Nfkb1*^-/-^ mice and to 25.1 cells (+/-1.6, *p*<.01) in *Nfkb2*^-/-^ mice (Figures 5A and B).

**Figure 5:**
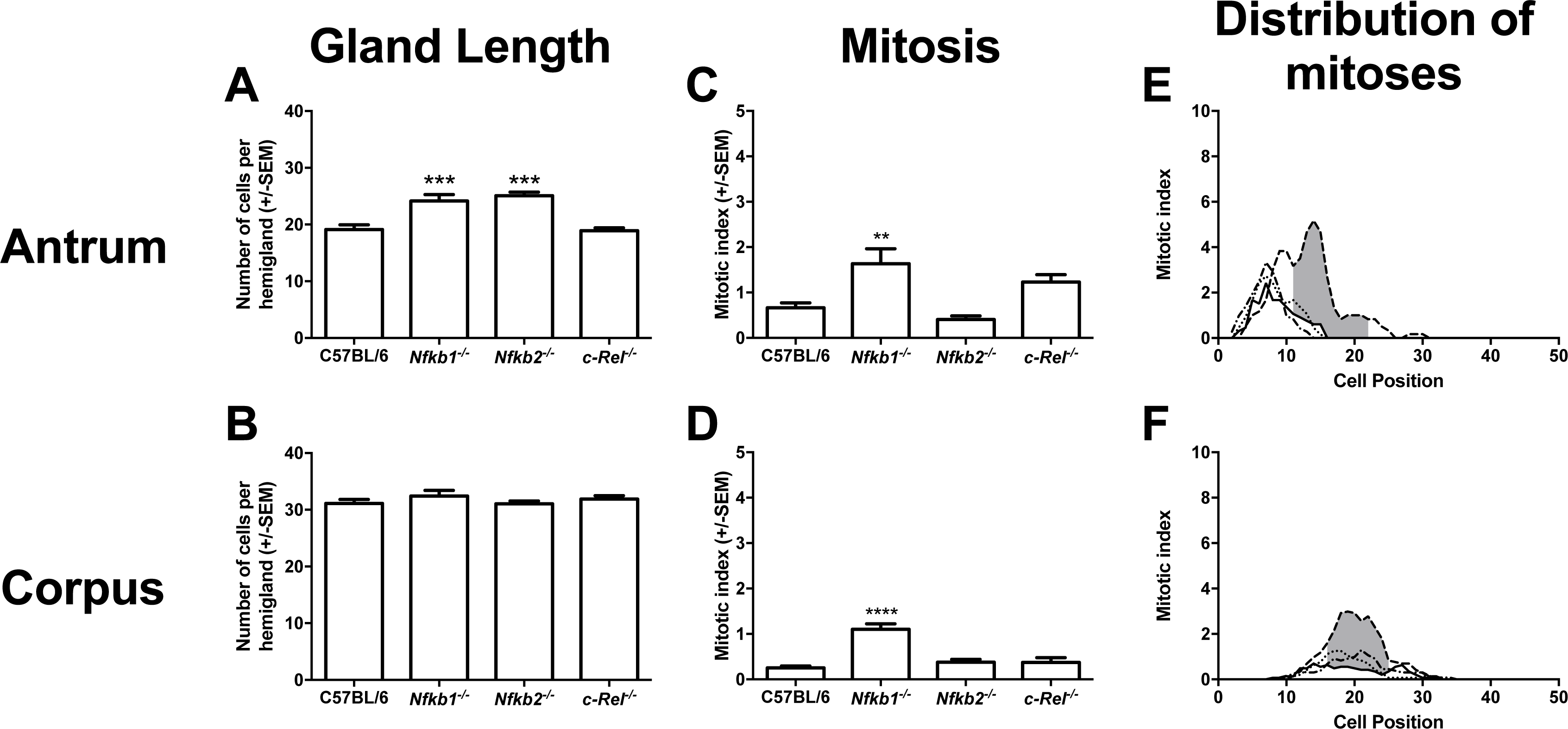
Impact of NF-κB subunit deletion on gastric gland length (A+B), number of gastric epithelial cells undergoing mitosis (C+D) and location of mitoses within the gastric gland (E+F) of untreated mice. A, C and E represent events in the gastric antral mucosa, B, D and F show data from the gastric corpus. Statistically significant differences tested by 1-way ANOVA and Dunnett’s test A-D, ** *p*<0.01, *** *p*<.001 **** *p*<0.0001. E and F: Solid line represents distribution of mitotic cells in WT mice, broken line *Nfkb1*^-/-^ mice. Shaded area marks region of gland where distribution of mitotic cells differs significantly in WT vs *Nfkbl*^-/-^ mice, *p*<0.05 by modified median test. N=6 for all experimental groups.

An average of 0.66%(+/-0.24) of antral gland cells were mitotic in untreated WT mice, and these were distributed between cell positions 3 and 15 with peak mitotic index at cell position 7. In *Nfkb1*^-/-^ mice the number of mitotic cells observed was higher (1.6 +/- 0.73%, *p*<0.05, Figure 5C), and the distribution of mitoses occurred over a wider area (cell positions 4-25), with the position of peak mitotic index shifted up the gland to cell position 14. Statistically significant increases in mitotic indices were observed at cell positions 11-22 in *Nfkb1*^-/-^ compared to WT mice (*p*<0.05, Figure 5E). No statistically significant differences in the number or distribution of mitotic cells were identified in the gastric mucosae of mice lacking NF-κB2 or c-Rel.

In gastric corpus mucosa, 0.25%(+/-0.10) of cells were mitotic in WT mice (Figure 5D). These events were distributed between cell positions 10 and 31, with peak mitotic index at cell position 14 (Figures 5D and F). In *Nfkb1*^-/-^ mice 1.1%(+/-0.29, *p*<.0001) of gastric corpus cells were mitotic and mitotic cells were distributed between cell positions 11 and 32, with peak mitotic index at cell position 19. Increased gastric corpus mitotic index (*p*<0.05) was observed in *Nfkb1*^-/-^ mice compared with WT mice between cell positions 16 and 25 (Figures 5D and F).

In untreated WT mice 0.17%(+/-0.25) of cells in the gastric antrum were identified as morphologically apoptotic; these events were distributed in low numbers from cell position 4-12. Neither the number nor the distribution of apoptotic cells was significantly different in the antrum of untreated *Nfkb1*^-/-^ mice (Figure 6A, C and E). In the corpus of untreated WT mice, 0.04%(+/-0.04) of cells were morphologically apoptotic, and were distributed between cell positions 8 and 26. By this measure, the number of apoptotic cells in the gastric corpus was increased 4.8-fold in untreated *Nfkb1*^-/-^ mice (0.19% +/-0.15, *p*<0.05), and the apoptotic cells were distributed over a marginally wider region of the corpus gland than in WT mice (cell positions 5-32) (Figure 6B, D and F).

**Figure 6:**
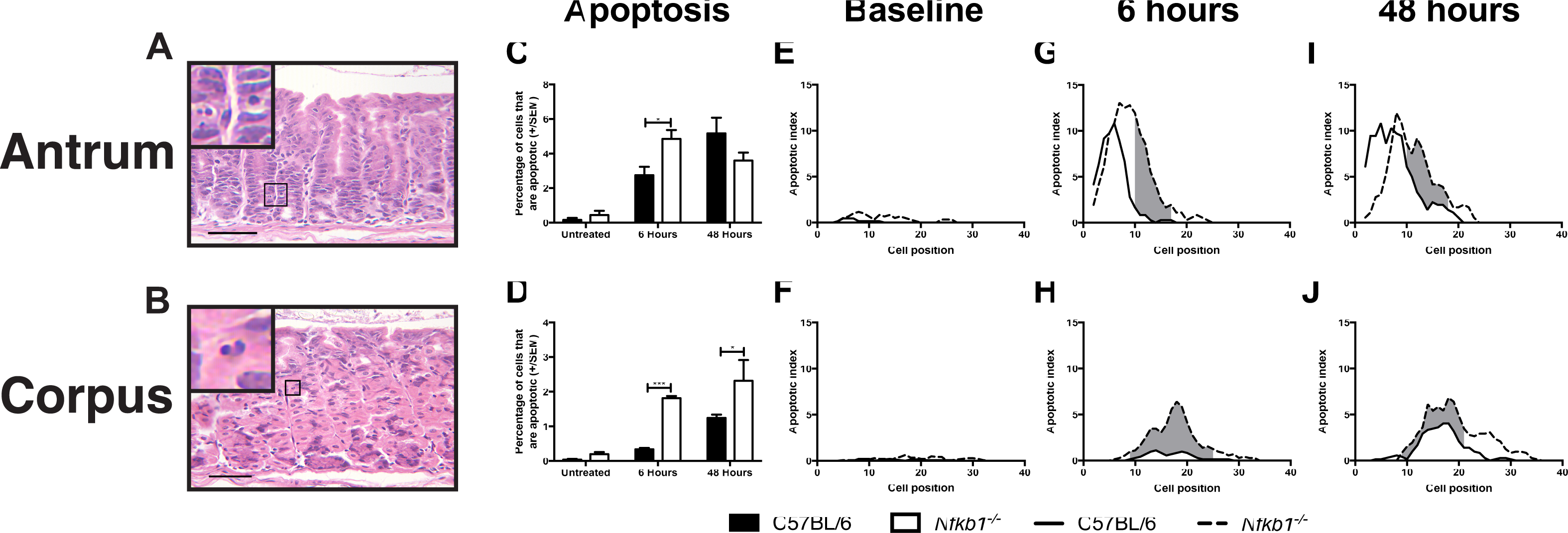
Impact of NF-κB1 deletion on gastric epithelial apoptosis in untreated and 12Gy γ-irradiated mice 6 and 48-hours after irradiation. A+B: Representative images of gastric corpus and antrum respectively stained with H+E, arrows highlight apoptotic epithelial cells. Scale bars 100μm. C+D: percentage of gastric epithelial cells undergoing apoptosis in gastric antrum and corpus respectively. Statistically significant differences tested by 2-way ANOVA and Dunnett’s test. * *p*<0.05, *** *p*<0.001. E-J: Apoptotic cells plotted by cell position in the gastric antrum (E, G and I) and corpus (F, H and J) of untreated and irradiated WT and *Nfkb1*^-/-^ mice. Solid lines represent distribution of cells in WT mice, broken lines *Nfkb1*^-/-^ mice. Shaded areas mark regions of glands where distribution of apoptotic cells differs significantly in WT vs *Nfkb1*^-/-^ mice, *p*<0.05 by modified median test. N=6 for all experimental groups.

γ-irradiation entirely suppressed mitosis in the gastric tissues of both WT and *Nfkb1*^-/-^ mice at both 6- and 48-hours post irradiation. However, six hours after γ-irradiation, the numbers of apoptotic cells identified in both the corpus and antrum of *Nfkb1*^-/-^ mice were higher than in similar tissues from WT mice (Figure 6C and D). In the antrum, 2.8%(+/-1.1) of gastric antral cells were apoptotic in WT mice, with apoptotic cells distributed between cell positions 2 and 17 and peak apoptotic index at cell position 6. Six hours after γ-irradiation, 4.4%(+/-1.5, p<0.05) of antral cells were apoptotic in *Nfkb1*^-/-^ mice, with apoptotic cells observed between cell positions 2 and 24, and peak apoptotic index at cell position 7. Significantly increased apoptosis was observed in *Nfkb1*^-/-^ mice between cell positions 10 and 17(*p*<0.05, Figure 6G).

Six hours after γ-irradiation 0.34%(+/-0.04) cells were apoptotic in the gastric corpus of WT mice, compared to 1.8%(+/-0.07) cells in *Nfkb1*^-/-^ mice (*p*<.0001). Apoptotic cells were distributed between cell positions 9 and 28 in WT mice, with peak apoptotic indices observed at cell positions 13 and 19. In *Nfkb1*^-/-^ mice, apoptosis was observed between cell positions 6 and 33, with peak apoptosis at cell position 18. Increased apoptosis was observed in the gastric corpus of *Nfkb1*^-/-^ mice between cell positions 9 and 25 (*p*<0.05, Figure 6H).

Forty-eight hours after γ-irradiation the differences in apoptotic cell number between WT and *Nfkb1*^-/-^ mice were less marked, particularly in the gastric antrum. At this time-point 5.2%(+/-2.0) of antral epithelial cells were apoptotic in WT mice compared to 3.6% of cells in *Nfkb1*^-/-^ mice (*p*=0.10, Figure 6C). Apoptosis was observed in WT mice between cell positions 2 and 20, with peak apoptosis at cell position 5. The distribution of apoptosis in *Nfkb1*^-/-^ mice was shifted slightly up the antral gland compared to WT mice, with apoptotic cells being observed between cell positions 2 and 23, and peak apoptosis at cell position 8. Increased apoptosis was observed in *Nfkb1*^-/-^ mice between cell positions 10 and 18 (*p*<.05, Figure 6I).

In the gastric corpus, 1.2%(+/-0.09) of cells were apoptotic in WT mice 48 hours after γ-irradiation, compared to 2.3%(+/-0.60) in *Nfkb1*^-/-^ mice (*p*<0.05 Figure 6D). At this time-point, apoptotic cells were observed between cell positions 4 and 30, with peak apoptosis at cell positions 17 and 18 in WT mice. In *Nfkb1*^-/-^ mice, apoptotic cells were observed between cell positions 9 and 35, with peak apoptosis at cell position 14. Increased apoptosis was observed between cell positions 9 and 21 (*p*<0.05, Figure 6J).

### Nfkb1^-/-^ animals have primed extrinsic pathway apoptosis mechanisms in the gastric epithelium

To investigate the mechanism underlying the increased gastric epithelial apoptosis in mice lacking *Nfkb1,* we extracted mRNA from mucosal samples of mice either without treatment, or 6 hours following γ-irradiation. We performed real-time PCR assays to quantify the expression of 8 regulators of apoptosis that have previously been shown to be under the transcriptional regulation of NF-κB signalling^15, 16, 17, 18, 19, 20, 21^. No statistically significant differences in the expression of the inhibitors of apoptosis c-IAP1, c-IAP2 or xIAP were identified (Figures 7A-C), nor were changes in the expression of p53, BCl-2 or BCL_XL_ observed (Figures 7D-F). In contrast, Fas and FasL expression were both upregulated (1.8- and 4.8-fold, p<.01 and p<0001 respectively, Figures 7G and H) in the gastric mucosa of untreated *Nfkb1*^-/-^ mice compared to WT mice. Six hours after γ-irradiation, Fas and FasL transcript abundance increased in WT mice to levels similar to those identified in *Nfkb1*^-/-^ mice at baseline, suggesting that this pathway is activated in response to γ-irradiation.

**Figure 7:**
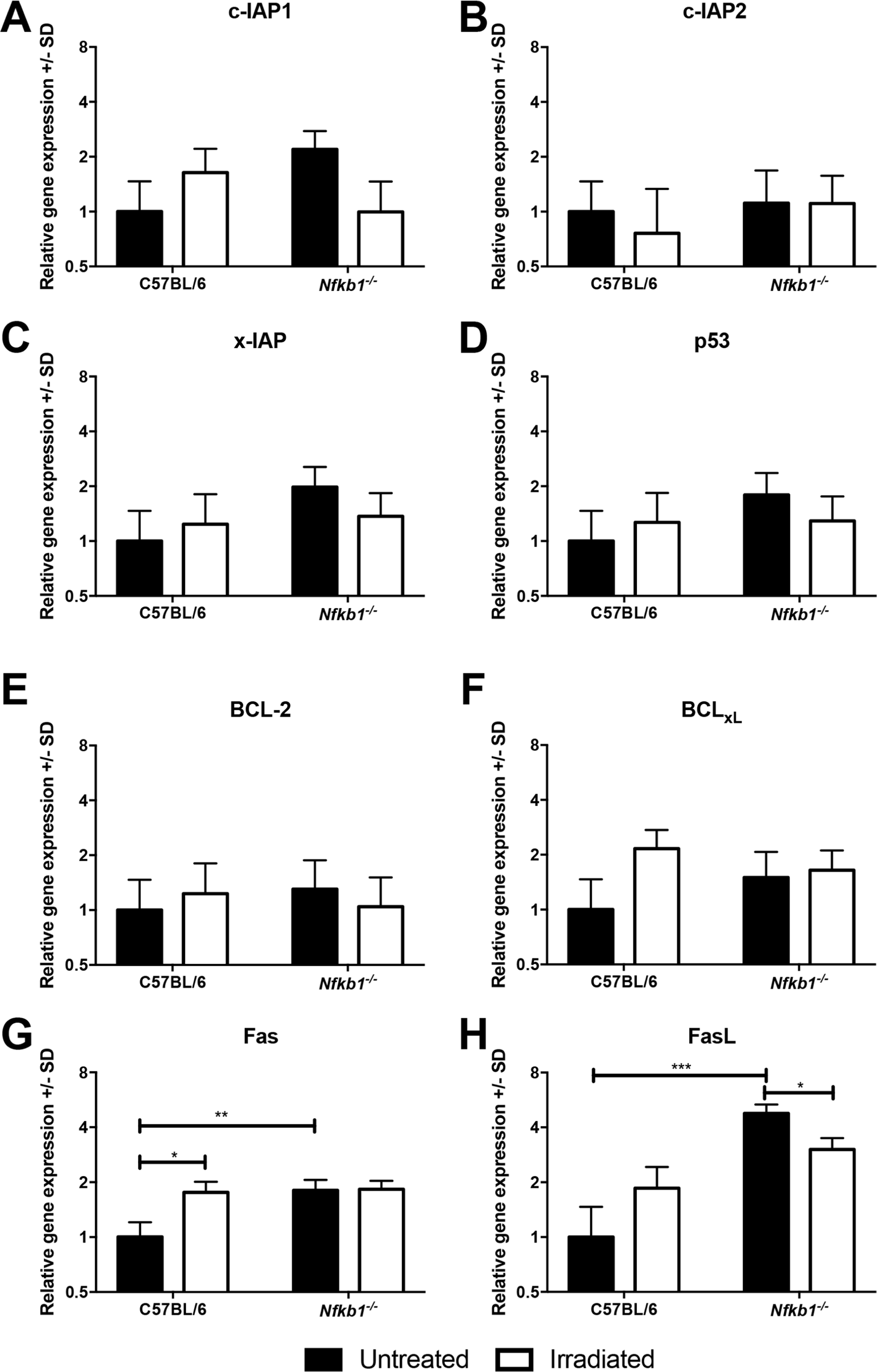
Relative gene expression of specified regulators of apoptosis under transcriptional regulation of NF-κB signaling. Filled bars represent untreated mice, open bars animals culled 6 hours after 12Gy γ-irradiated. Mean and standard deviation of Log_2_ transformed relative abundance for each transcript are plotted. Statistically significant differences tested by 2-way ANOVA and Dunnett’s test, * *p*<0.05, ** *p*<0.01, *** *p*<.001. N=3 for all experimental groups.

## Discussion

These data demonstrate that NF-κB signaling is involved in regulating the development of gastric epithelial metaplasia and atrophy following tamoxifen administration. In this model the most significant NF-κB subunit appears to be NF-κB1, as mice lacking NF-κB2 and c-Rel demonstrated few differences in response compared to WT mice. This contrasts with previous observations from our own laboratory that have demonstrated differential regulation of *H. felis* induced gastric preneoplastic pathology in mice lacking NF-κB2 as well as NF-κB1.

The differences in gastric epithelial pathology observed in *Nfkb1*^-/-^ mice following tamoxifen administration were associated with a more pronounced proliferative response and increased epithelial cell apoptosis. This increase in proliferation was observed throughout the gland. This is consistent with Burclaff *et al’s* recent reports that describe the development of a metaplastic progenitor source following tamoxifen induced dedifferentiation of chief cells^22^.

Further investigation of the DNA damage responses of *Nfkb1*^-/-^ mice demonstrated an increased sensitivity to γ-irradiation induced epithelial cell apoptosis than WT mice. This suggests that the difference in response to tamoxifen may reflect a more generalized difference in DNA damage responses in these mice, rather than a specific tamoxifen related event. This is supported by earlier studies which have shown that following γ-irradiation, *Nfkb1*^-/-^ mice exhibited more small intestinal apoptosis than WT mice^23^.

The dynamics of NF-κB and p53 mediated signaling in response to DNA damage have been subject to systematic modelling in recent years^24^. This work has demonstrated that these mechanisms are closely linked, and form a complex regulatory network for DNA damage responses. It is therefore unsurprising that abrogation of NF-κB signaling pathways leads to altered DNA damage responses. Nonetheless, it is striking that following tamoxifen administration this effect was almost exclusively associated with NF-κB1 deletion.

We adopted a candidate gene approach to try to identify specific targets of NF-κB signaling that could explain the observed differences in apoptosis. Whilst there are compromises associated with this approach, we identified potential priming of the CD95/FasL pathway in untreated mice lacking NF-κB1. Whilst this pathway is not the most frequently studied in DNA damage induced apoptosis, there are both *in-vitro*^25^ and *in-vivo*^26^ studies that demonstrate its role in modulating radiation induced apoptosis. Further studies that address the functional effect of components of this pathway on tamoxifen induced gastric lesions may provide further insight into this mechanism.

Previous data have demonstrated that gastric epithelial cells express the estrogen receptor, and hence a direct abrogation of signaling through this receptor is a biologically plausible mechanism for inducing gastric atrophy, however, the similarity of lesions induced in the stomach following tamoxifen with that induced by known protonophores including DMP-777 has also promoted the concept of tamoxifen acting as a direct epithelial cell toxin^10^. Our DNA damage assays suggest that genotoxic stress may contribute to the gastric lesions induced by tamoxifen. However further studies investigating the underlying mechanism of tamoxifen induced gastric murine pathology are required.

We conclude that signaling involving NF-κB1 regulates gastric epithelial pathology in response to a second model of gastric atrophy and metaplasia. As in *H. felis* infection, following tamoxifen administration, signaling mediated by NF-κB1 appears to suppress gastric epithelial cell turnover, and DNA damage responses. This is coupled with altered baseline expression of Fas and FasL in mice lacking NF-κB1, which may contribute to excess apoptosis following genotoxic stress. Given that these animals demonstrate differences in susceptibility to both genotoxic stress, and susceptibility to gastric pre-neoplasia, further studies investigating how abrogation of NF-κB signaling pathways influences chemical carcinogens induced gastric carcinogenesis would provide interesting insight into the importance of these pathways during gastric carcinogenesis.

In the current study, we observed no differences between wild-type and *Nfkb2*^-/-^ mice. This is in contrast to *H. felis* infection, following which *Nfkb2*^-/-^ mice were almost entirely protected from gastric epithelial pathology. This contrast suggests that the effects of *Nfkb1*^-/-^ and *Nfkb2*^-/-^ deletion on gastric epithelial pathology are mediated through different mechanisms.

## Materials and Methods

### Mice

All murine procedures were performed at the University of Liverpool Biomedical Services Unit in a specific pathogen free research facility under appropriate UK Home Office licensing, and following University of Liverpool Research Ethics Committee approval.

Animals were maintained with a standard 12-hour light/dark cycle and received standard rodent chow and water *ad-libitum* throughout the experimental procedures. Female C57BL/6 wild-type (WT) mice were purchased from Charles River (Margate, UK) and maintained for a minimum 7-day acclimatization prior to use in experiments. *Nfkb1*^-/-^, *Nfkb2*^-/-^ and *c-Ref*^-/-^ mice (as previously described^27^) were bred and maintained on a C57BL/6 genetic background at the University of Liverpool.

### Animal procedures

Tamoxifen (Cayman Chemicals, Cambridge Biosciences, Cambridge, UK) was prepared as previously described in ethanol and corn oil^8^. Groups of at least 5 female mice aged 10-12 weeks were administered either 150mg/kg tamoxifen, or vehicle via a single intra-peritoneal injection. 72 hours later animals were euthanized by cervical dislocation, and gastric tissues were harvested for histology.

Whole body γ-irradiation was performed by exposure to a Caesium-137 source in a GammaCell closed source irradiator. Groups of at least 6 female mice aged 10-12 weeks were exposed to a single 12Gy fraction of γ-irradiation. Animals were returned to standard housing conditions prior to being euthanized at 6 or 48 hours post procedure by cervical dislocation. Gastric tissues were harvested for histology. In a separate experiment, similar groups of 3 mice were treated identically prior to cervical dislocation at 6 hours. From these animals, the luminal surface of the stomach was scraped to generate gastric mucosa enriched samples and flash-frozen in liquid nitrogen prior to nucleic acid extraction.

### Immunohistochemistry

Standard immunohistochemical techniques were adopted throughout. Heat induced epitope retrieval was performed for all antigens in sodium citrate buffer, pH 6.0. Primary antibodies used were rabbit anti-H^+^/K^+^ATPase (SantaCruz SC-84304), rabbit anti-Ki67 (AB16667, AbCam, Cambridge, UK), rabbit anti-γH2AX (#9718, Cell Signalling, New England Biolabs, Hertfordshire, UK). Secondary detection of all antibodies was performed using IMMpress anti-Rabbit polymer (Vector Laboratories, Peterborough, UK) and SigmaFast 3,3’-diaminobenzidine (DAB) (Sigma-Aldrich, Dorset, UK).

### Quantitative histology

Quantitative histology was performed using previously validated cell positional scoring systems for H+E stained and immunohistochemically stained tissues. Briefly, well oriented gastric glands were identified and individual cells were scored morphologically, or based on positive immunostaining, starting with cell position 1 at the base of the gland, and extending up the gland to the luminal surface. Each gland unit scored from base to apex of the gland is described as a hemigland, for each gastric corpus section a total of 30 hemiglands were scored^28^. Visual analogue scoring of gastric preneoplastic lesions was performed as previously described by Rogers *et al.*^11^ Cleaved-caspase-3 immunostaining was quantified based on the number of positively stained cells per high powered field. For this score, 10 non-overlapping fields of gastric mucosa were visualized using a x40 objective per tissue section. The number of positively stained cells per high power field was counted, and mean score per section calculated.

### Quantitative Real-Time PCR

Total RNA was extracted and reverse transcribed using the Roche Highpure RNA Tissue kit and Transcriptor reverse transcription kits respectively. Real-Time PCR was performed on a Roche LightCycler 488 instrument, and assays were designed using the Roche Universal Probe Library. Details of primers, probes and amplicons are included in table 1. All real-time PCR reagents were sourced from Roche UK, Burgess Hill, UK.

### Statistics

Statistical analyses were performed using GraphPad Prism 7. Data were analyzed with 2-tailed, 2-way Student’s *t*-test, or 2-way analysis of variance (ANOVA) and Dunnett’s *post-hoc* analyses as appropriate. Differences in cell positional distributions were assessed using a modified median test as previously described^29^.

## Acknowledgements

We would like to thank Dr Jorge Caamano and Bristol Myers Squibb for donating the *Nfkb2*^-/-^ mouse colony, and Dr Jorge Caamano for providing the colonies of *Nfkb1*^-/-^ and *c-Rel*^-/-^ mice. MDB was funded by a CORE / British Society of Gastroenterology Development Grant and Wellcome Trust / University of Liverpool Institutional Strategic Support Fund grant under grant agreement number: 097826/Z/11/Z. MDB and DMP were supported by a North West Cancer Research project grant. MDB, JMW, RH and DMP were supported by the European Council’s Seventh Framework Programme (FP7/2007-2013) under grant agreement number 305564 (SysmedIBD).

